# Altered host pyruvate metabolism fuels and regulates fungal asexual reproduction

**DOI:** 10.1101/2024.08.22.609210

**Authors:** Mi Yeon Lee, Johan Jaenisch, Amanda G. McRae, Chihiro Hirai-Adachi, Katherine Louie, Leslie P. Silva, Benjamin P. Bowen, Trent R. Northen, Mary C. Wildermuth

## Abstract

Powdery mildew (PM) fungi are widespread pathogens impacting agricultural productivity. As obligate biotrophs, PMs get all nutrients from their host plant. Concurrent with asexual reproduction, the PM *Golovinomyces orontii* induces host lipid body accumulation in plant mesophyll cells underlying the epidermal cell that houses the fungal feeding structure. Herein, we show a PM-induced shift in plant primary metabolism, to bypass the pyruvate dehydrogenase complex (PDHc), acts in a dose-dependent manner to support host storage lipid production for PM acquisition. Not only are plant PDHc bypass-derived lipids incorporated into spore storage lipids, but they also act in a regulatory capacity to control the number of fungal reproductive structures per colony, ensuring the fitness of spores that are formed. Evolutionarily distant plant and animal obligate biotrophs manipulate host lipid metabolism for their own acquisition and we propose altered host pyruvate metabolism, bypassing the host PDHc bottleneck, as a common strategy to increase host source flux to acetate/acetyl-CoA for obligate microbial lipid acquisition.

## INTRODUCTION

Powdery mildew fungi infect a wide variety of crops and have the greatest impact on world agricultural productivity of any pest^1^. Furthermore, they are among the most rapidly-spreading pests^2^. As obligate plant biotrophs, powdery mildew (PM) fungi can only grow and reproduce on living plants. They alter plant cellular architecture and metabolism to acquire all nutrients from the plant, while limiting plant defense. The PM *Golovinomyces orontii* MGH1 (*Gor*) infects *Arabidopsis thaliana*, the resource-rich model plant dicot^3,4^. *Gor* infection of an *Arabidopsis* leaf begins with germination and differentiation of the spore, penetration of the plant cell wall and establishment of a fungal feeding structure (haustorium) in the host epidermal cell by 1 day post inoculation (dpi). The fungus then proliferates forming an extensive surface hyphal network. At 5 dpi, the mesophyll cells directly underlying the haustorium have undergone endoreduplication, resulting in increased DNA copy number and increased size^5,6^, and fungal asexual reproduction becomes apparent (**Fig. 1a**). These asexual reproductive structures, conidiophores (cp), carry chains of newly formed spores (conidia) filled with lipid bodies containing triacylglycerols (TGs) (**Fig. 1a-c**). Concurrently, lipid bodies accumulate in the host mesophyll cells underlying the epidermal cell housing the haustorium, as well as in the haustorium^7^. PM spore lipid composition is altered when plant lipid composition is artificially altered indicating PMs incorporate host lipids into spores^8^. Therefore, the reproductive fitness of the PM is defined by its ability to acquire these host lipid bodies. In addition, the lipid stores dictate spore competence when colonizing a new host as subsequent catabolism of spore storage lipids is associated with spore germination and differentiation^9^ and silencing PM TG catabolism genes limits proliferation^10^.

**Figure 1.**
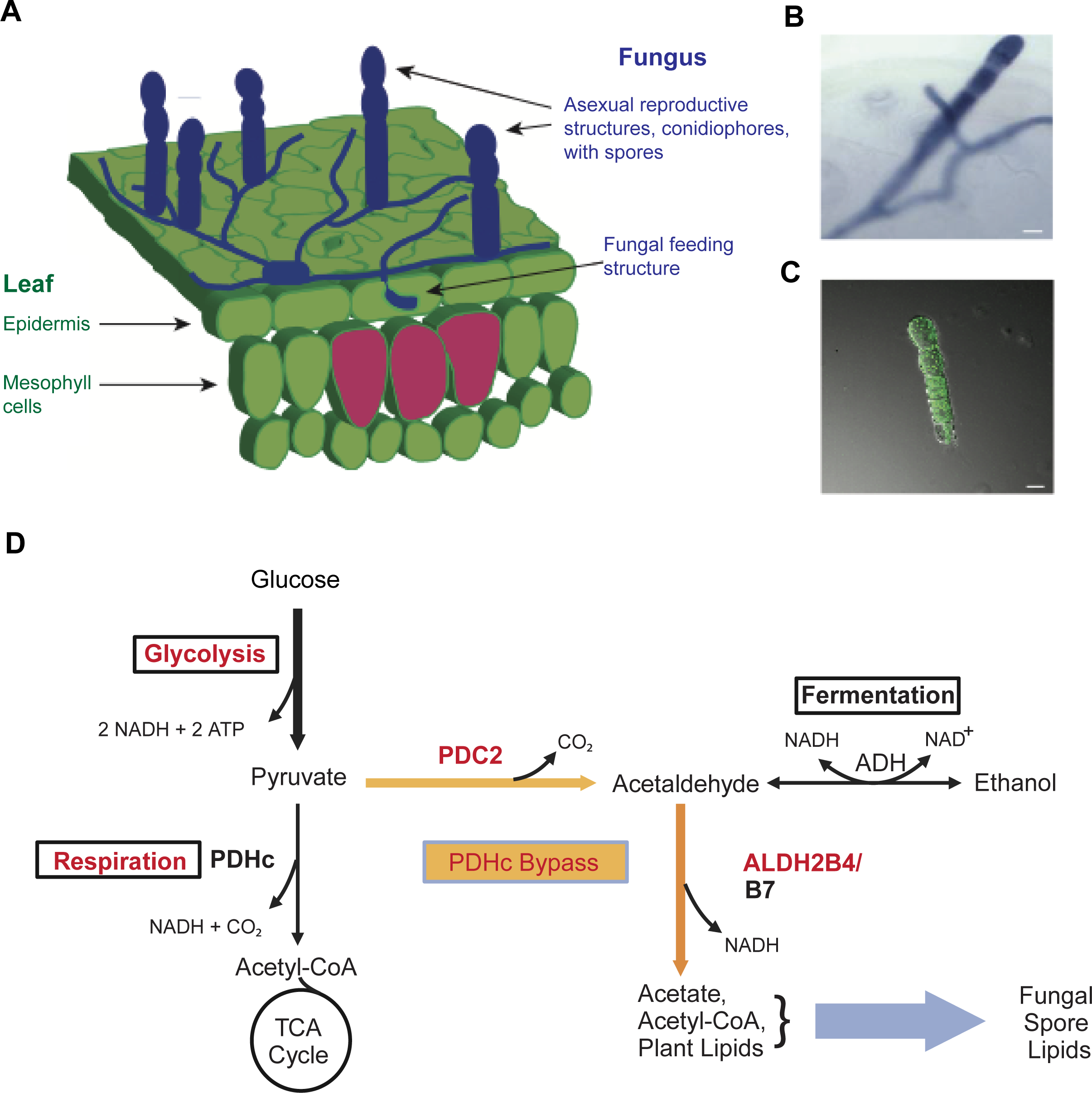
Powdery mildew manipulation of *Arabidopsis* primary metabolism at the infection site concurrent with its asexual reproduction. **A,** Side view depiction of PM infection of *Arabidopsis* leaf at 5 dpi. Magenta cells exhibit PM-induced endoreduplication^5^. **B-C,** Microscopic visualization of cps, the asexual reproductive structures, stained with trypan blue (**B**) or BODIPY 505/515 neutral lipid fluorescent dye (**C**); bar=20 µm. **D,** Simplified view of *Arabidopsis* primary metabolism highlighting use of the PDHc bypass for host lipid production and fungal lipid acquisition. Statistically significant enhanced expression at 5 dpi^5,15^ of GO processes (black box) and genes (red font).

How does the powdery mildew manipulate host primary metabolism to obtain the lipids it requires for asexual reproduction? With PM infection, leaf carbohydrates are mobilized to the fungal infection site for fungal acquisition^11–14^. As shown in **Figure 1d**, PM infection site transcriptome profiling at 5 dpi indicates enhanced glycolysis and respiration consistent with a localized increase in acetyl-CoA and energy production^5^. Surprisingly, the fermentation gene ontology (GO) term is also enhanced^5,15^. Fermentation utilizes a cytosolic pyruvate decarboxylase (PDC, EC 4.1.1.1) and alcohol dehydrogenase (ADH, EC 1.1.1.1) resulting in the conversion of pyruvate to ethanol **(Fig. 1d**). Alternatively, the fermentation GO term could reflect use of the pyruvate dehydrogenase complex (PDHc) bypass, as it is not separately specified. The PDHc bypass also employs a cytosolic PDC, but then converts acetaldehyde to acetate via an acetaldehyde dehydrogenase (ALDH; EC 1.2.1) (**Fig. 1d**).

Mature plant leaves do not typically utilize fermentation or the PDHc bypass. Fermentation is typically observed and studied during hypoxia (e.g. in hypoxic roots) when mitochondrial respiration is limited by oxygen availability. However, fermentation and/or use of the PDHc bypass has been observed in aerobic tissue during periods of high metabolic/energy demand, such as pollen tube or seedling development^16–18^. Aerobic fermentation could facilitate enhanced metabolic capacity by regenerating NAD+ for continued glycolysis and respiration^19^. In addition, use of the PDHc bypass could further increase acetyl-CoA and lipid production when glycolytic and respiratory capacities are saturated^18^. Below, we investigate the potential contributions of fermentation and the PDHc bypass to PM asexual reproduction. We further propose a localized induced shift in host primary metabolism to bypass the PDHc bottleneck as a shared strategy to support increased source flux to host lipids for use by plant and animal obligate biotrophs.

## RESULTS

### Host PDC2 supports PM asexual reproduction in a dose-dependent manner

As shown in **Figure 1d**, both fermentation and the PDHc bypass utilize cytosolic pyruvate decarboxylase. Of the four *A. thaliana PDC* genes, we focused on *AtPDC2* (At5g54960) as it exhibits enhanced expression in response to PM infection at 5 dpi^5^ and is most similar to petunia PDC2 (**Fig. S1a**) shown to play a role in the energetically demanding process of pollen tube elongation^16^. The *Arabidopsis pdc2-1* mutant (*pdc2*) contains a T-DNA insertion in the 5’UTR of the *PDC2* gene, resulting in a dramatic reduction in PM-induced *PDC2* expression at 5 dpi (**Fig. S1b-c**). We find *pdc2* plants have a visible decrease in PM growth and reproduction, with significantly less visible leaf coverage (**Fig. 2a**). Furthermore, microscopic analysis of PM colony development (cp/colony) shows a 41% reduction in cps formed for PM that develop on *pdc2* compared to wild type (WT) plants (**Fig. 2b**). Complementation of *pdc2* with *PDC2* restores cp/colony to WT levels (**Table S1**). Decreased PM asexual reproduction is not associated with dwarfism (**Fig. 2a)** nor cell death in *pdc2* (**Fig. S1d**).

**Figure 2.**
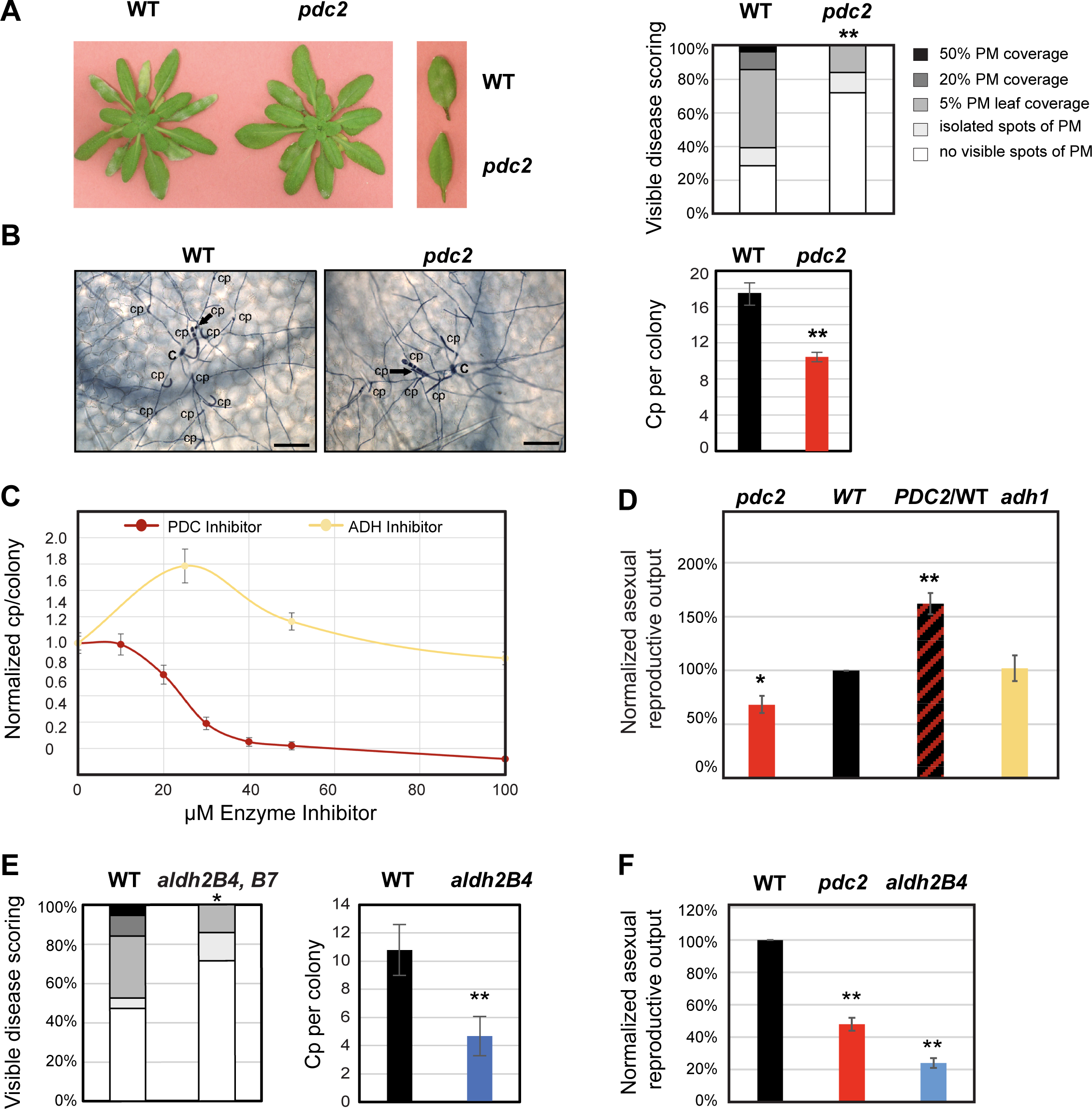
Host PDHc bypass not fermentation supports PM asexual reproduction. **A**, PM-infected whole plants and representative leaves of WT and *pdc2* at 10 dpi. Visible disease scoring comparing WT and mutant; n=20-25 plants per genotype. **B,** Microscopic analysis of leaves harvested at 5 dpi and stained with trypan blue for analysis of cp/colony; C= initial germinated conidia; cp= conidiophore; Bar= 100 µm. n≥30 colonies per genotype. **C**, Impact of chemical inhibition of Arabidopsis PDC or ADH enzymes on cp/colony. Fully expanded mature *Arabidopsis* leaves 2 dpi were detached and petioles placed in ½ MS agar plates containing enzyme inhibitors. Five days later, cp/colony were assessed as in **B**. Mean with SE is shown; n=45 colonies per concentration per genotype. **D,** PM spore production (spores/mgFW leaf) is normalized to values on WT plants infected in parallel. *PDC2/WT* is homozygous plant line with an extra copy of *PDC2* driven by its native promoter stably introduced into WT. **E, (left)** Visible disease scoring comparing WT and *aldh2b4,b7 (aldh2B)* mutant as in **A**; **(right)** cp/colony for WT and *aldh2B* leaves harvested at 5 dpi as in **B**. **F,** Normalized PM spore production (as in **D**). For all experiments, mean is shown with SE for one experiment. Independent experiments gave similar results. Statistical significance used Student’s t-test or Pearson’s Chi-squared analysis. *= *p*-value ≤0.05; **= *p*-value ≤0.001.

To explore the impact of PDC dosage on PM asexual reproduction, we developed a detached leaf system that allows us to supply biochemical inhibitors through the *Arabidopsis* leaf petiole. Fully expanded mature leaves from 4 week-old plants inoculated with PM are detached at 2 dpi when the PM feeding structure has formed, but asexual reproduction has not yet occurred. The petioles are placed in agarose with varying concentrations of inhibitor and cp/colony are assessed 5 days later. We find host PDC inhibition reduces cp/colony to <20% of uninhibited levels (**Fig. 2c**). Furthermore, the impact of the PDC chemical inhibitor on fungal asexual reproduction is dose-dependent with a minimal inhibitory concentration (MIC)_50_ of 27 µM for the experiment shown and a mean MIC_50_ of 34±4 µM (Mean±SE, n=3 independent experiments). This MIC_50_ is similar to that found for the bovine parasite *Tritrichomonas foetus* when its growth is dependent on PDC and to the IC_50_ of the purified PDC enzyme^20^. We then employed a genetic approach to examine whether PM asexual reproduction correlates with *PDC2* dosage. We generated WT plants with an additional copy of *PDC2* under control of its native promoter and assessed spore production per mg leaf fresh weight for *pdc2*, WT, and homozygous WT lines with an extra copy of *PDC2 (PDC2/WT)*. Spore production decreases by 45% on *pdc2* compared to WT plants (**Fig. 2d**), similar to the percent reduction in cp/colony **(Fig. 2b**). And, fungal spore production increases proportionally with *PDC2* dosage (**Fig. 2d**).

### Host PDHc bypass, not fermentation, supports PM asexual reproduction

To explore whether host fermentation is used to support PM asexual reproduction (**Fig. 1d**), we inhibited ADH function using our detached leaf biochemical inhibitor assay. PM asexual reproduction is not reduced by ADH chemical inhibition, instead the number of cp/colony is enhanced **(Fig. 2c**). Arabidopsis *ADH1* (At1g77120) is not induced in response to PM at 5 dpi^5,6^. Furthermore, an *adh1* null mutant shows no difference in spore production (**Fig. 2d)** confirming that the fermentation pathway does not support PM asexual reproduction.

The *Arabidopsis* PDHc bypass utilizes acetaldehyde dehydrogenases ALDH2B4 (At3g48000) and ALDH2B7 (At1g23800) to convert acetaldehyde to acetate for lipid synthesis^18^. *ALDH2B4* expression is enhanced in response to PM at 5 dpi, while *ALDH2B7* expression remains very low^5^. Because the *aldh2B4,B7* double null mutant (*aldhB2*), disrupted in both genes, was previously shown to be required for a strong PDHc bypass-associated phenotype^18^, we first employed it in our studies. Similar to *pdc2*, the *aldhB2* double mutant supports less visible PM growth compared to WT (**Fig. 2e, Fig. S2a**), with fewer cp/colony (53% reduction compared to WT). Decreased PM asexual reproduction is not associated with cell death in *aldh2B* and the plants exhibit no pleiotropic phenotypes (**Fig. S2**). Because *ALDH2B4* is induced at the PM infection site at 5 dpi^5^, we also assessed PM asexual reproduction on the *aldh2B4-1* (*aldh2B4*) single null mutant. We found spore production is reduced 76% on *aldh2B4* compared to WT plants (**Fig. 2f**) indicating a dominant role for ALDH2B4 in this process. Together, these results show the Arabidopsis PDHc bypass, using PDC2 and ALDH2B4, operates in a dose-dependent manner to support powdery mildew asexual reproduction.

### Host PDHc bypass supplies lipid precursors for powdery mildew spores

In order to determine whether C derived from the plant PDHc-bypass pathway is incorporated into newly formed PM spore TGs, we used an experimental setup similar to our biochemical inhibition experiments and tracked incorporation of supplied ^13^C-ethanol into lipids of the detached washed leaf and newly formed PM spores that develop on the leaf (**Fig. 3a**). Because only the ADH enzymatic reaction is reversible, supplied ^13^C-ethanol would be utilized by the induced PDHc bypass (**Fig. 1d**). Wei et al. (2009) successfully used this approach to show contribution of the *Arabidopsis* PDHc bypass to FA synthesis in seedlings^18^. It should be noted that supplied ethanol is provided via the leaf petiole to the leaf and therefore not directly accessible to the PM. Furthermore, *Gor* does not contain an *ADH* gene (**Excel File S1**).

**Figure 3.**
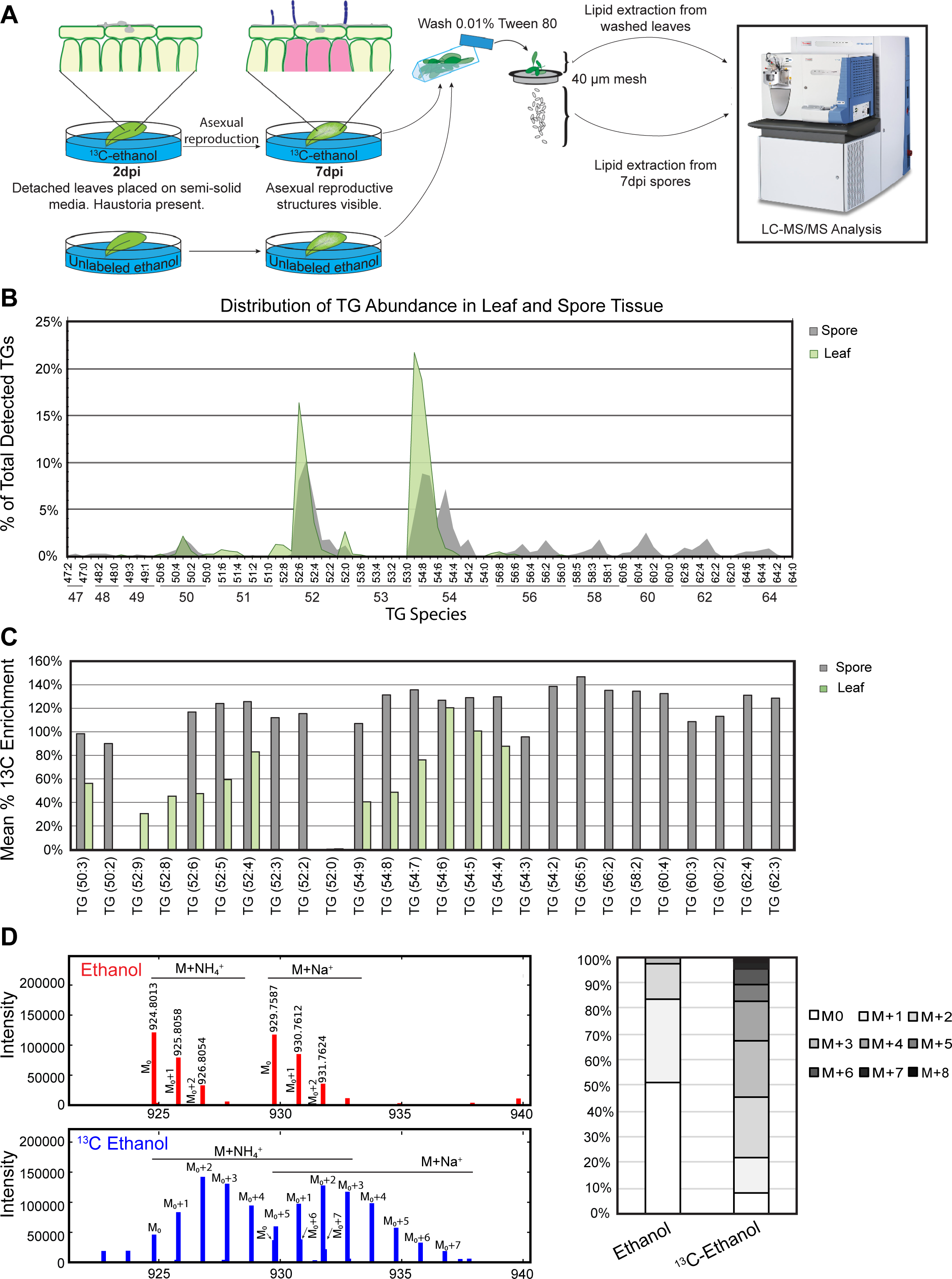
Incorporation of plant PDHc outputs into leaf and spore triacylglycerols. **A,** Overview of experimental design. Mature fully expanded Arabidopsis leaves 2 dpi are detached and petioles are placed in agar containing either ^13^C-ethanol or unlabeled ethanol. Five days later, leaves are removed and lipids are extracted from fungal spore and washed leaf samples for analysis by LC-MS/MS. **B,** Relative abundance of TGs in leaf and spore tissue is calculated based on the value of unlabeled mean M0 peak height per mg fresh weight (FW) tissue; n=3. **C,** Percent ^13^C enrichment for TG species extracted from ^13^C-ethanol-fed vs ethanol-fed leaves, calculated as (^13^C-ethanol-fed M_0+1_/M_0_ - ethanol-fed M_0+1_/M_0_) / (ethanol-fed M_0+_1/M_0_), shown for TG species representing ≥1% total TG tissue abundance. **D,** MS1 spectra for spore TG(56:6) from spores isolated from infected leaves fed ethanol (top) or ^13^C-ethanol (bottom) with isotopologue analysis showing relative contribution of each isotopologue to total (sum of isotopologues); shown for NH_4_^+^ adducts. Independent experiments gave similar results.

We found that while TG species of infected leaves are highly abundant in PM spores that develop on those leaves, spore TGs do not simply parallel infected leaf TG profiles (**Fig. 3b, Excel File S2**). TGs containing very long chain fatty acids (VLCFAs) (56:x to 64:x) are more abundant in spores, representing 24% of total spore TGs by abundance, while infected leaf TGs contain only 56:x TGs comprising 1% of total infected leaf TG abundance. MS2 analysis of a subset of the spore TGs confirms the presence of C20-C24 acyl chains (e.g. **Fig. S3).** In addition, spore TGs tend to have acyl chains that are slightly more saturated than leaf TGs (**Fig. 3b, Fig. S4, Excel File S2**). Infected leaf TGs exhibit statistically significant M_0+1_/M_0_ enrichment (**Fig. 3c**, **Excel File S2**), reflecting efficient ^13^C-ethanol/ethanol uptake by the leaf and incorporation into lipids. Spore TGs are also ^13^C enriched, with percent enrichments higher than that of the associated infected leaf TGs (**Fig 3c, Excel File S2**). For example, for the two most abundant TG classes (52:x and 54:x) which together comprise 93% of leaf TGs and 69% of spore TGs, spore TG ^13^C enrichment is on average 1.8-fold greater than infected leaf TG enrichment. Furthermore, spore TGs containing VLCFAs display high levels of enrichment, averaging 131%. ^13^C enrichment calculations are based on M_0+1_/M_0_ to enhance specificity but thereby underestimate total ^13^C enrichment as we see multiple label incorporation into both leaf and spore TGs (**Fig. S4**). For example, spore TG 56:6 MS spectra and associated isotopologue analysis show multiple labels incorporated into TG 56:6 of spores that develop on ^13^C-ethanol supplied leaves vs. unlabeled ethanol-fed leaves (**Fig. 3d**). Incorporating all isotopologues above M_0_ in the enrichment analysis results in 1127% enrichment for this spore TG sample compared to 176% enrichment using M_0+1_/M_0_ **(Excel File S2).** Moreover, spore TG 56:6 isotopologues ≥ M_0+3_ represent 55% of total isotopologues for spores that develop on ^13^C-ethanol fed leaves compared to 3% for unlabeled ethanol fed leaves (**Fig. 3d**).

### PDHc bypass output controls the number of cp/colony, but not their development or fitness

To investigate whether plant PDHc bypass-derived products control cp development once initiated, we microscopically evaluated cp development. *Gor* cp development consists of formation of a basal foot cell, followed by a generative cell that successively elongates and divides to produce a conidium (spore) in its upper section (**Fig. 1b**). This process is repeated resulting in a chain of conidia with the oldest at the tip. Microscopic analysis of cp development by stage shows it is not impacted by chemical inhibition of PDC (at the IC_50_) or by development on *pdc2* plants **(Fig. 4a-b**). These results indicate cp initiation but not subsequent development is dependent on *Arabidopsis* PDHc bypass function.

**Figure 4.**
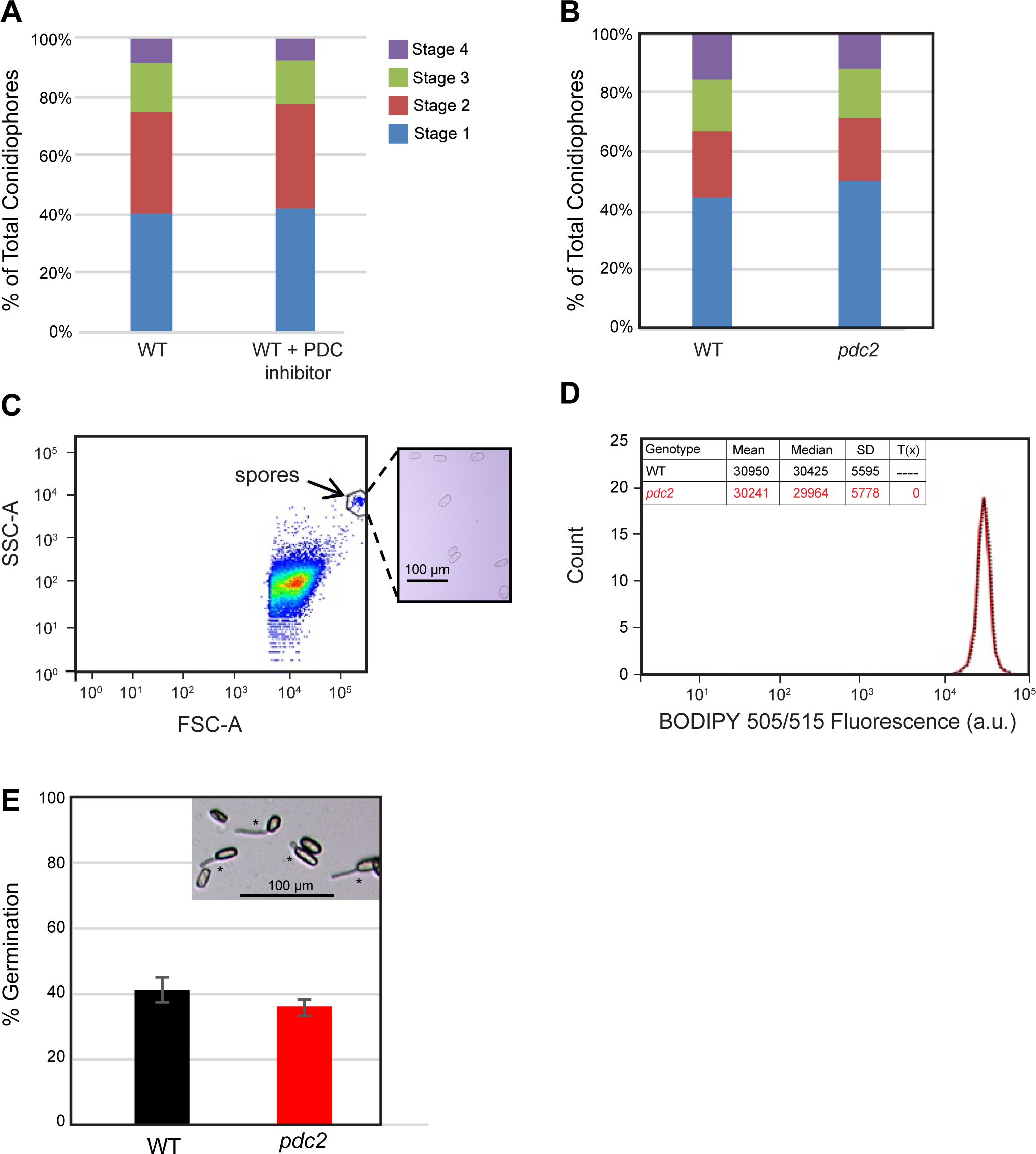
PDHc bypass output regulates the number of cp/colony but not their development or fitness. **A-B**, Mean percent of total cp at each developmental stage at 5 dpi. Stage 1(basal foot cell) to Stage 4, containing progressively more conidia. PM developed on (**A**) WT leaves with and without PDC inhibitor at the MIC_50_ or (**B**) WT compared with *pdc2* leaves. Data shown is for one independent experiment (cp from 45 colonies, n=167-629). Pearson’s Chi-squared analysis found no statistical difference. An independent experiment gave similar results. **C**, Spores from PM that developed on WT and *pdc2* plants were stained with neutral lipid dye, BODIPY 505/515, and fluorescence was measured using a flow cytometer. Spore suspension was visualized by forward scatter (FSC-A) and side scatter (SSC-A) to identify spores (gated). Inset shows spores sorted by a BD Influx Sorter to verify gating methods. **D**, The fluorescence of 200 (±10%) spores developed on WT (black) and *pdc2* (red) was measured. Fluorescence intensity distributions were compared using Chi-Squared T(x) statistical analysis. Analysis shows no significant difference (T(x)=0). Two additional independent experiments show a similar result. **E,** The germination rate (mean %, ± SE) of spores that developed on WT and *pdc2* plants was determined (3 samples, each with n≥100). Student’s t-test of the germination rates shows no significant difference (p≥0.05). Two additional independent experiments show a similar result. Inset shows spores in germination assay; * indicates germinated spore.

To ascertain whether the lipid content of spores that do form are impacted by reduced plant PDHc bypass function, we used flow cytometry analysis of spores coupled with BODIPY 505/515 neutral lipid staining. We find no difference in spore lipid content for PM that develop on WT compared with *pdc2* plants (**Fig. 4c-d**). Qualitative microscopic comparisons also show no difference in spore lipid body content or morphology. To assess spore fitness, we ascertained whether the germination rate of PM spores formed on *pdc2* plants differs from those that develop on WT plants. No difference in spore germination rates is detected (**Fig. 4e**).

## DISCUSSION

### Plant PDHc bypass output both regulates and provides precursors for powdery mildew asexual reproduction

Asexual reproduction by *G. orontii* at 5+ dpi involves the formation of cps that contain a chain of spores each filled with lipid bodies containing TGs (**Fig. 1**). At 5+ dpi, lipid bodies filled with TGs are observed in mesophyll cells at the infection site and in the fungal haustorium^7^. Site specific profiling of *Gor*-infected leaves at 5 dpi revealed the expression of genes associated with glycolysis, respiration, and fermentation to be significantly enhanced^5,15^. Herein, we show that a shift in plant primary metabolism to use the PDHc bypass, not fermentation, acts in a dose-dependent manner to support PM asexual reproduction (**Figs. 1d, 2**). Furthermore, using ^13^C substrate labeling we *directly* show that fungal spore TGs incorporate host-derived lipids (**Fig. 3**). Consistent with PDHc bypass function (to produce acetate/acetyl-CoA), ^13^C-label supplied via the plant PDHc bypass results in ^13^C enrichment of TGs in infected leaves and the spores that develop on these leaves **(Fig. 3, Excel File S2, Fig. S4**). Spore TGs exhibit on average 1.7-fold higher ^13^C M_0+1_/M_0_ enrichment than their infected washed leaf TG counterparts. This indicates spore development is fueled by localized infection-site specific metabolic changes not observed across the entire leaf (**Fig. 1**).

*Gor* spore TG composition reflects infected leaf TG composition, but with increased VLCFAs and an overall slight increase in saturation (**Fig. 3, Excel File S2, Fig. S4)**. VLCFAs, C20-C24, have been reported to be highly prevalent in PM conidia^21–23^ as we observe for *Gor* TGs. Spore TGs containing VLCFAs not present in PM-infected leaves exhibit high ^13^C enrichment (**Fig. 3; Excel File S2**) indicative of remodeling of acquired host lipids by the PM. Moreover, transcriptional profiling of *Gor^10^* and other PMs (e.g. *E. necator^24^*) over the course of infection found FA desaturases and elongases that exhibit dramatically increased expression concurrent with asexual reproduction.

Not only does the Arabidopsis PDHc bypass supply lipid precursors for PM acquisition during asexual reproduction, but its activity also controls the extent of asexual reproduction. The development of colonies is constrained by PDHc bypass activity with AtPDC1 dose-dependency shown using both genetic and biochemical inhibition approaches (**Fig. 2**). The cp that are initiated develop normally, forming spores replete with lipid bodies (**Fig. 4**). As spore storage lipids decrease with germination and fungal TG lipases are expressed at this time, catabolism of spore storage lipids likely provides nutrients/energy for spore germination and differentiation, prior to formation of the fungal feeding structure^9,25^. Silencing *Gor* TG lipase reduced PM proliferation, consistent with this hypothesis^10^. Therefore, we ascertained whether the germination rate of PM spores formed on *pdc2* plants differs from those that develop on WT plants and found they germinate equally well (**Fig. 4**). It has long been known that lipids can act as developmental signals in fungi^26-27^. Our results suggest a host PDHc-derived product acts as such a PM sporulation signal or precursor to a sporulation signal, linking the extent of initiation of asexual reproductive structures to host metabolic status.

### Powdery-mildew induced flux to TGs: addressing the source term

Plant oils are important commodities; therefore enhancing TG production has received much attention^28,29^. It is clear that increasing TG biosynthetic enzymes alone do not sufficiently increase oil yield and that a multi-targeted approach is required. Metabolic control analysis (MCA) provides a framework to address and identify important components that contribute to flux through a metabolic pathway^30,31^, with flux control coefficients quantifying the impact that altering the activity of an enzyme or group of enzymes (e.g. in a control block) has on the flux of the metabolic pathway. A recent detailed MCA of TG formation in oilseed rape identified three blocks of reactions controlling flux to seed TGs^32^. Control over flux to TGs is dominated by source block 1 that generates acetyl-CoA and has a calculated flux control coefficient of 0.68 out of 1 total. In contrast, the downstream sink blocks 2 and 3 have less control over flux to TGs with calculated flux control coefficients of only 0.1 and 0.22, respectively.

At the PM infection site, increased glucose, from starch breakdown and sugar transport to the new sink site, provides elevated substrate for glycolysis. Glycolysis is enhanced and pyruvate, the product of glycolysis, accumulates beyond the capacity of the PDHc to utilize it, even with increased expression of genes involved in respiration^5,15^. The Arabidopsis PDHc bypass genes which are preferentially induced at the infection site concomitant with PM asexual reproduction^5^ then function to convert this pyruvate surplus to acetate/acetyl-CoA for lipid production (**Figs. 1d, 2-3**).

Much of the reason the PDHc bypass is so critical to increased flux to acetyl-CoA and derived products is due to the biochemical properties of cytosolic PDC. The Arabidopsis PDHc E1 operates at µM concentrations of pyruvate (S_0.5_ ∼ 45 µM^33^) compared to S*_0.5_* of 3-18 mM for cytosolic AtPDC2^34^ allowing AtPDC2 to make use of highly elevated pyruvate. Furthermore, AtPDC2 exhibits substrate cooperativity, similar to yeast PDC^35^. Finally, with saturation of the PDHc, cytosolic pyruvate accumulation rapidly acidifies the cytosol, further activating AtPDC2^34^. AtPDC2 exhibits a pH optimum of 5.7, with a 7-fold increase in *K_cat_* to 20.4 s^-1^ and a 3.7-fold increase in *V_max_* to 18.38 µmole/min mole protein at pH 5.7 compared to pH 7.5^34^. Taken together, our data explains how flux through the PDHc bypass (source block 1 to acetyl-CoA) acts as a dominant contributor to TG production. Consistent with MCA predictions^32^, a null mutant in the Arabidopsis TG biosynthetic enzyme diacylglycerol transferase 3 (DGAT3) responsible for PM-induced TGs that support spore production (control block 3) supports 30% less PM asexual reproduction^7^, while limitation of PDHc bypass enzyme activity (control block 1) through genetic disruption or biochemical inhibition reduces PM spore production by as much as 80% in a dose-dependent manner (**Fig. 2**). Importantly, reduction in PM asexual reproduction for the PDHc bypass mutants is not associated with activated plant defense as indicated by cell death or reduced size (**Figs. 2, S1-S2**). By contrast, mutants in core plant FA biosynthesis enzymes KAR1, KAS I, and FATB exhibit cell death and reduced size, complicating interpretation of their impact on PM spore production^8^.

### Shared strategies to increase host lipid availability for use by obligate biotrophs

It has long been known that host lipid manipulation and acquisition by obligate biotrophic pathogens and symbionts of animals supports and regulates demanding phases of their life cycles. Intracellular bacterial pathogens of mammals such as *Mycobacterium tuberculosis (Mbt)*^36^, mammalian *Apicomplexa* intracellular parasites such as *Toxoplasma gondii* and *Plasmodium falciparum*^37, 38^, and viruses including human cytomegalovirus (CMV) ^39^ utilize induced host lipid bodies^40,41,42^ and/or specific host lipid species. For example, *Mbt* colonization of the lung involves the manipulation of local host cell machinery to enhance FA and cholesterol production for its own acquisition and gene regulation^43^. Whereas, human malarial parasite *P. falciparum* uses host (lyso)phosphatidylcholine to fuel and regulate its sexual differentiation^38,44^.

Our understanding of plant obligate biotroph manipulation of and reliance on host lipids is more recent. It has been best studied for arbuscular mycorrhizal fungi (AMF), endosymbionts that colonize roots of a broad array of plant species^45,46^. AMF direct host lipid flux to 16:0 2-monoacylglycerol or a derivative which appears to be the dominant plant lipid acquired by AMF^47,48,49^. Furthermore, *in vitro* experiments with AMF suggest a host-derived lipid regulates AMF reproduction^46^, as we see for the PM (**Figs. 2, 4**). Similar to AMF, the protist *Plasmodiophora brassicae,* the causative agent of clubroot on rapeseed and other cruciferous plants, is a FA auxotroph, dependent on host lipids^50^. By contrast, PMs have the capacity to synthesize FAs; however, genomic evidence supports a preference for host lipid acquisition^51^. Herein, we experimentally show this preference for host lipids during the metabolically and energetically demanding process of PM asexual reproduction (when extensive lipids are required for storage in newly formed spores); limitation of host precursor supply to TGs dramatically reduces spore production (**Figs. 2,3**). Furthermore, the PM manipulates host metabolism to direct host lipid flux to TGs, which appear to be the dominant lipid acquired by the fungus (Figs 2,3; and ^7^).

Understandably, initial efforts to define the lipid dependency of obligate biotrophs on their host sought to define the dominant host lipid(s) acquired by the obligate biotroph and how host FA synthesis and lipid assembly (sink blocks 2 and 3 in the MCA framework of Fell et al. (2023)) are manipulated to increase the production of this lipid. Elucidating source controls resulting in increased acetyl-CoA (block 1) has largely focused on controls over the localized increase in glucose observed for these systems. How does increased glucose then result in increased acetyl-CoA driving flux to lipids?

Herein, we show PM manipulates plant primary metabolism to increase flux to TGs by increasing source control to acetate/acetyl-CoA through use of the non-respiratory plant PDHc bypass (**Fig. 1d**). Plant PDHc bypass genes also exhibit enhanced expression at the AMF-plant host interaction site^15,52^, suggesting the host PDHc bypass may also be utilized to support AMF lipid acquisition. Similarly, *P. brassicae* likely makes use of the host PDHc bypass to increase host lipid supply as a study focused on host fermentative metabolism and its impact on *P. brassicae* growth and reproduction found that the *Arabidopsis pdc2* mutant, but not the *adh1* mutant, resulted in reduced clubroot symptoms and *P. brassicae* spore formation^53^; *aldh* mutants were not tested. Therefore, a shift in plant primary metabolism to use the PDHc bypass may be a common means by which obligate plant biotrophs increase source supply to lipids for their own gain. In yeast, the PDHc bypass has long been part of efforts to engineer increased production of biotechnology products derived from acetyl-CoA including TGs^54,55,56^ supporting the importance of this source pathway to increased lipid flux.

While animals don’t have the plant PDHc bypass genes PDC and ALDH2B, other means by which they can bypass the host PDHc bottleneck to increase acetate/acetyl-CoA and lipid production when glucose is prevalent are being uncovered^57,58^. In the context of obligate biotrophy, human CMV was found to redirect host metabolism to convert glucose to acetate and acetyl-CoA for lipogenesis and viral growth^59^. Furthermore, *Plasmodium* hepatocyte infection rates are positively correlated with enhanced glycolysis, non-respiratory acetyl-CoA formation, and cholesterol biosynthesis^60,61^. In addition, altered pyruvate metabolism that bypasses the PDHc bottleneck to increase acetyl-CoA and lipids is a hallmark of certain cancers^62^. Together, these studies suggest that manipulation of eukaryotic host primary metabolism to bypass the PDHc bottleneck is a shared strategy of diverse obligate biotrophs to increase host acetate/acetyl-CoA and lipid synthesis for their own gain.

## Acknowledgements

This work was supported by National Science Foundation grants IOS-0958100 to MCW and MCB-1617020 to MCW and TRN. We thank Dr. Hans van Veen (Utrecht University, Netherlands) for assistance with statistical analyses of distribution data, Hang Xue (UC Berkeley, UCB) for assistance with figures, and Hang Xue and Claudine Tahmin (UCB) for critical review of the manuscript.

## Author contributions

MYL, CH, JJ, and AGM performed plant molecular genetic and powdery mildew analyses. JJ also performed ^13^C feeding experiments and lipid extractions; LC-MS/MS lipid analyses were done by LS, KL, and BB under the direction of TRN. AGM also assessed spore fitness. MCW directed the research and wrote the manuscript with review and input from all authors.

## STAR METHODS

### Plants

All *Arabidopsis thaliana* mutants are in the Col-0 background except for the *aldh2B4-1,aldh2B7-1* double mutant which is in the Ws-0 background. The appropriate WT control is used in all experiments. Homozygous *pdc2-1* (SALK_066678C), *aldh2B4-2* (SALK_078568), and *adh1-1* (SALK_066824) T-DNA insertion lines (Alonso et al. 2003), were obtained from the Arabidopsis Biological Resource Center. *aldh2B4-1/B7-1* double mutant and *aldh2B4-2* lines were previously characterized as null mutants (Wei et al. 2009) as was *adh1-1* (Hwang et al., 2011). **Figure S1** provides more information on the *pdc2-1* mutant. All mutants were confirmed by PCR with primers in **Table S2**.

To generate stable transgenic plants, constructs were introduced into *Agrobacterium* and transformed into WT Col-0 or *pdc2-1* by floral dip (Clough and Bent, 1998). The 2523 bp product containing the *PDC2* promoter (*PDC2pro*) and *PDC2* gene was amplified from *A. thaliana* Col-0 genomic DNA and introduced into pCAMBIA 1300 in frame with previously introduced enhanced GFP (from pEGFP-N1, Clonetech, CA) and *CAMV* 35S terminator site. All PCR products were verified by DNA sequencing. The plasmid containing *PDC2pro::PDC2-eGFP* (below) was introduced into *Agrobacterium tumefaciens* strain GV3101 and transformed into *Arabidopsis pdc2-1* and WT. Lines with insertions were selected by resistance to Hygromycin and GFP expression, and inserted sequences were verified. Primers are provided in **Table S2**. Homozygous stable transgenic lines with one introduced copy of *PDC2* in *pdc2 (PDC2pro::PDC2-eGFP/pdc2-1*) referred to as *PDC2/pdc2* were used for the complementation experiment (**Table S1**) while lines with one introduced copy of *PDC2* in WT (*PDC2pro::PDC2-eGFP/*WT) referred to as *PDC2/*WT was used for the dosage experiment, shown in **Figure 2**. Plant lines generated in this study are available from the corresponding author M. Wildermuth.

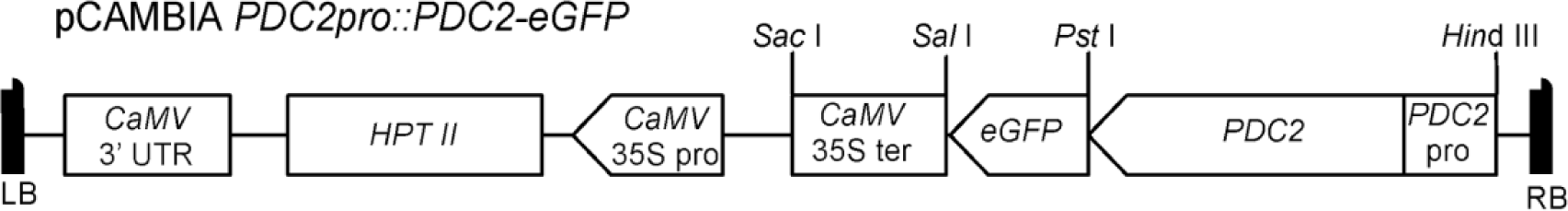

Schematic diagram of pCAMBIA 1300-derived plasmid with inserted PDC2pro*::PDC2-eGFP.* LB, left border; RB, right border; *CaMV,* Cauliflower Mosaic Virus*; HPT II,* hygromycin phosphotransferase gene; pro, promoter; ter, terminator.

### Powdery mildew assays

*A. thaliana* WT and mutants were grown, evenly spaced, using an alternating pattern with 12 plants per box in Percival AR66L environmental chambers at 22°C, 70% RH, and 12 h photoperiod with photosynthetically active radiation (PAR) of 120-150 µmol m^-2^ s^-1^. At 4 weeks, each box was infected with conidia from 2 half-covered leaves (10-14 dpi) of *G. orontii* MGH1 isolate (Plotnikova et al., 1998) using a uniform settling tower method with mesh screen as in Chandran et al., 2009. Visual and microscopic PM disease phenotypes were performed as described in Chandran et al. 2009, with visual disease phenotypes scored at 10-11 dpi based on percent visible coverage of the leaf. 20-25 plants were evaluated per genotype for one experiment. For microscopic analysis, three fully expanded leaves per plant (leaves ∼7-9 post emergence) were excised at 5 dpi, cleared and stained with trypan blue for visualization of fungus and counting of asexual reproductive structures (conidiophores) per colony, or cell death. Leaves from four plants per genotype were analyzed with at least 30 PM colonies assessed. Conidiophore developmental stage was noted at the time of assessment, and ranged from Stage 1 (initial nub of the basal foot cell is visible microscopically) to Stage 4 containing progressively more conidia. Independent experiments gave similar results. For assessing differences in distributions (visual disease score and cp development) distributions, Pearson’s Chi-squared test with Yates continuity correction was utilized.

For powdery mildew spore counting, the procedure of Wessling et al. (2014) was optimized to enhance reproducibility. At 8-10 dpi following a moderate to heavy inoculation dose (3-4 fully covered leaves), three fully expanded mature leaves per plant for 5-6 plants of each genotype in a box were excised, weighed, and vortexed with 15 ml 0.01% Tween-80 (Sigma). Following leaf removal, the solution was filtered through a 40 µm mesh (Sysmex) and conidia in the filtrate were pelleted at 4,000 x *g* for 5 min. Pellets were resuspended in 200-600 ul final volume ddH_2_0 and 10 ul aliquots used in an IMPROVED NEUBAUER 0.1mm deep Brightline Houser Scientific hemacytometer. All 9 fields were independently counted and the mean and standard deviation (stdev) calculated to ensure sample counts were reproducible. In addition, a lower threshold of 80,000 spores/gFW for WT was utilized to improve reproducibility. Note that each planted box is an independent biological sample directly comparing WT and mutant plant lines. 4-6 independent biological samples were assessed per mutant vs WT comparison. Student’s t-test was utilized for statistical analysis of paired comparisons (e.g. WT vs. mutant cp/colony or spores/mgFW).

For detached leaf experiments, 4-week old WT plants were grown in environmental chambers and infected with a moderate dose of *Gor* conidia using settling towers, as above. At 2 dpi, three fully expanded leaves were excised and the petioles were placed in semi-solid media (1/2 MS salts, 0.8% BactoAgar, with either inhibitor or ethanol) in 100 mm x100 mm x15 mm square petri dishes, sealed with micropore tape (3M), with 12 leaves per dish for inhibitor experiment and 20 leaves per dish for ethanol feeding experiment. The petri dishes were then placed back in the environmental chamber.

For the biochemical inhibition studies, the PDC inhibitor omeprazole (Sigma-Aldrich) or the ADH inhibitor pyrazole (Sigma-Aldrich) was dissolved in DMSO to make a working stock solution and added to the agar media solution just before pouring. Volumes added were adjusted with DMSO for consistency across all concentrations including no inhibitor. The series of concentrations were assessed on separate plates in the same experiment. Five days later (7 dpi), leaves were removed from agar and stained for microscopic counting of cp/colony as above. At least 45 colonies were counted per inhibitor dose, with on average 5 colonies observed from each of 9 leaves. Independent experiments gave similar results.

For the ethanol feeding experiment, 10 mM ^13^C-ethanol (Sigma-Aldrich) or ethanol was added to the agar media just before pouring. Five days later (7 dpi), leaves were removed from the agar and washed in 0.01% Tween-80 (Sigma). The resulting solution was filtered through 40µm mesh (Corning), spun down at 4,000 x *g* for 5 minutes to pellet the spores, resuspended in 500µL ddH_2_0 and spun down. The pelleted spores were weighed, frozen using liquid Nitrogen, finely ground using a TissueLyserII (Qiagen) with 3 mm size balls, and immediately extracted for lipid analysis (below). Three biological replicates per experiment, with ∼50 infected leaves per replicate resulting in ∼ 0.17 gFW spores per sample. For the washed leaves, a subset of the washed pool of leaves of ∼ 0.075 gFW leaf material was used per sample, ball milled as above and immediately extracted for lipid analysis (below). For one independent experiment, three spore samples and three washed leaf samples from both control ethanol and ^13^C-ethanol plates were analyzed. An independent repeat of the entire experiment confirmed findings. Note that *G. orontii MGH1* has no bonafide PDC or ADH candidate (**Excel File S1**).

### Lipid LC-MS/MS analyses

Lipids were extracted from spore and washed leaf samples from ethanol or ^13^C-ethanol agar plates using a modified Bligh & Dyer extraction (1:1:0.9 methanol:chloroform:H_2_0) (Li-Beisson et al. 2010). 300µL of chloroform phase was recovered per leaf or spore sample, and dried under nitrogen. The dried extracts were re-suspended in 200 µL of 3:3:4 IPA:ACN:MeOH, and run immediately. A triacylglyceride (TG) internal standard mix (LM6000, Avanti) was used to ensure retention time reproducibility. Samples were run on an Agilent 1290 (Agilent Technologies, Santa Clara, CA) UHPLC connected to a QExactive mass spectrometer (Thermo Fisher Scientific, San Jose, CA) with the following chromatographic method, in both positive and negative mode. Lipids were run on a reversed phase 50mm x 2.1 mm, 1.8 µm Zorbax RRHD (Rapid Resolution High Definition) C18 column (Agilent Technologies) with a 21 min gradient and 0.4 mL/min flow rate, with 2 µL injections. The mobile phases used were A: 60:40 H_2_O:ACN with 5mM ammonium acetate, 0.1% formic acid, and B: 90:10 IPA:ACN with 5mM ammonium acetate (0.2% H_2_O), 0.1% formic acid. The system was held at 20% B for 1.5 min, followed by an increase to 55% B over 2.5 min, and a subsequent increase to 80% B over 6 min. The system was then held at 80% B for 2 min, before being flushed out with 100% B for 5 min, and re-equilibrated at 20% B over 5 min. The QExactive parameters were as follows: MS resolution was set to 70,000, and data was collected in centroid mode from 80-1200 m/z. MS/MS data was collected at a resolution of 17,500 with a collision energy step gradient of 10, 20, and 30. All solvents were HPLC grade unless specified.

Lipid Data Analysis: Lipids, focused on TGs, were identified by comparing retention times and MS/MS fragmentation patterns in positive ion mode to standards either run in-house, the NIST 11 Mass Spectral Library (Kind et al., 2009), or from several online databases, including LipidMaps (Fahy et al., 2007) and Metlin (Smith et al. 2005). Calculations used peak height of NH_4_ ^+^ adducts, run in positive ion mode. Relative abundance = unlabeled mean M_0_ peak height/mg leaf fresh weight (FW), with n=3. M_0_ peak heights of <1E05 were considered to be below reliable detection threshold. A two-tailed T-test (unpaired with equal variance) was used to assess statistical significance between treatments; n=3 per treatment. Percent ^13^C enrichment = (Mean/mgFW ^13^C-ethanol supplied M_0_+1/M_0_ – Mean/mgFW unlabeled ethanol supplied M_0_+1/M_0_)/Mean/mgFW unlabeled M_0_+1/M_0_). Percent label incorporation = (M_0_+1 + M_0_+2*2 + M_0_+3*3 ….)/((M_0_ + M_0_+1 + M_0_+2 + M_0_+3…)*# of mass isotopologues), calculated as in Doomun et al., 2016. See **Excel File S2** for TG identification and data processing with representative calculations. Data is available at the EMBL-EBI Metabolights data repository (Yurekten et al., 2023), with ID www.ebi.ac.uk/metabolights/MTBLS1114.

### Powdery mildew spore lipid analysis using flow cytometry

As above, *Arabidopsis* plants are grown in a Percival using an alternating pattern of WT and mutant genotypes within each box to ensure equal growing conditions and even inoculation. At 4 weeks, *A. thaliana* is settling tower-inoculated with a moderate inoculum of *Gor* MGH1 per box. Nine days later, three fully expanded mature leaves per plant are harvested from 9 plants per genotype in the same box. To liberate spores, leaves are vortexed in 0.01% Tween 80 in ddH_2_0 for 30 seconds. Spore suspensions are filtered through 40 μm mesh and centrifuged at 4000 x *g*. The neutral lipid dye BODIPY 505/515 (ThermoFisher Scientific D3921) at a final concentration of 1μg/mL is added to spore suspensions which are then analyzed using LSR Fortessa X20 flow cytometer. Samples are interrogated with a 488 nm laser and signal captured with a FITC (505LP 530/30) filter. Sample acquisition uses BD FACSDiva Software (v7.0) with subsequent gating of spores via FlowJo (v10.1); see Figure 4. Each biological replicate measures the fluorescence intensity of 200 (±10%) spores shed from WT or mutant *pdc2-1* plants within the same box. Fluorescence intensity distributions are compared using Chi-Squared T(x) statistical analysis in FlowJo v.10 software. Two additional independent experiments were performed with a similar result.

### Powdery mildew spore germination assay

At 11 dpi, three mature fully expanded leaves from WT and *pdc2-1* plants are harvested and tapped over a glass slide to collect spores. Leaves from three plants per genotype are utilized. Slides are stacked in a petri dish with 3 ml ddH_2_O water to ensure high relative humidity, closed and covered with aluminum foil for 1 hr. Petri dishes are then exposed to PAR ∼100 uEm^-2^s^-1^ for 5 hr. At least 100 spores were observed per slide, with three slides per genotype. A Student’s t-Test (Two-Tailed,Two-Sample Assuming Equal Variance) is used to compare germination rates between genotypes. Two additional independent experiments showed similar results.

## SUPPLEMENTARY INFORMATION OVERVIEW

- **Supplemental Document 1:** Supplemental Overview, Figures S1-S4, Tables S1-S2.
- **Excel File S1**
- **Excel Workbook File S2**

**Figure S1.** Characterization of Arabidopsis *pdc2-1* T-DNA insertion mutant, supports Figure 2 and STAR Methods.

**Figure S2.** Representative images of *aldh2B4,B7* double mutant compared with WT plants infected with powdery mildew, supports Figure 2.

**Figure S3.** Spore TGs contain C20-C24 acyl chains (MS2 analysis), supports Figure 3.

**Figure S4.** Representative TG signatures for spore and washed leaf lipids for detached leaves supplied with ethanol vs. ^13^C-ethanol, supports Figure 3.

**Table S1:** Complementation of *pdc2* mutant, supports Figure 2.

**Table S2:** Primers Used, supports STAR Methods.

**Excel File S1.** Reciprocal tBLASTn results show no PDC or ADH candidate in *G. orontii MGH1*, supports STAR Methods. Provided as a separate file.

**Excel Workbook File S2.** TG identification, relative abundance and ^13^C enrichment results, supports Figure 3. Provided as a separate file.

## Methods References

Alonso, et al. (2003) Genome-wide insertional mutagenesis of *Arabidopsis thaliana*. Science 301:653–657.

Chandran D., Tai Y.C., Hather G., Dewdney J., Denoux C., Burgess D.G., Ausubel F.M., Speed T.P., Wildermuth M.C. (2009) Temporal global expression data reveal known and novel salicylate-impacted processes and regulators mediating powdery mildew growth and reproduction on Arabidopsis. Plant Physiol. 149:1435–51.

Clough S.J. & Bent A.F. (1998) Floral dip: a simplified method for Agrobacterium-mediated transformation of Arabidopsis thaliana. Plant J. 16:735–743.

Doomun N.E., Loke S., O’Callaghan S., Callahan D.L. (2016) A simple method for measuring carbon-12 fatty acid enrichment in major lipid classes of microalgae using GC-MS. Metabolites 6:42.

Fahy E., Sud M., Cotter D., Subramaniam S. (2007) LIPID MAPS online tools for lipid research. Nucleic Acids Research 35:W606–W612.

Hwang J.H., Lee, M.O., Choy, Y.-H., Ha-Lee, Y.-M., Hong C.B., Lee, D.-H. (2011) Expression profile analysis of hypoxia responses in Arabidopsis roots and shoots. Journal of Plant Biology 54:373.

Kind T., Wohlgemuth G., Lee D.Y., Lu Y., Palazoglu M., Shahbaz S., Fiehn O. (2009). FiehnLib: mass spectral and retention index libraries for metabolomics based on quadrupole and time-of-flight gas chromatography/mass spectrometry. Analytical Chemistry 81:10038–10048.

Li-Beisson Y. et al. (2010) Acyl-Lipid Metabolism, The Arabidopsis Book 2010:e0133.

Plotnikova J.M., Reuber T.L., Ausubel F.M. & Pfister D.H. (1998) Powdery mildew pathogenesis of Arabidopsis thaliana, Mycologia, 90:6, 1009–1016; DOI:10.1080/00275514.1998.12026999

Smith C.A., Maille G.O., Want E.J., Qin C., Trauger S.A., Brandon T.R., Custodio D.E., Abagyan R., Siuzdak G. (2005) Metlin: a metabolite mass spectral database. Therapeutic Drug Monitoring 27:747–751.

Wei Y., Lin M., Oliver D.J., Schnable P.S. (2009) The roles of aldehyde dehydrogenases (ALDHs) in the PDH bypass of Arabidopsis. BMC Biochemistry 10:7.

Weßling R. & Panstruga, R. (2014) Rapid quantification of plant-powdery mildew interactions by qPCR and conidiospore counts. Plant Methods 8:35.

Yurekten O., et al. (2023) MetaboLights: open data repository for metabolomics. Nucleic Acids Research, 2023, gkad1045, doi:10.1093/nar/gkad1045, PMID:37971328.

## REFERENCES

1. Glawe, D.A. (2008). The powdery mildews: a review of the world’s most familiar (yet poorly known) plant pathogens. Annu. Rev. Phytopathol. 46, 27–51. 10.1146/annurev.phyto.46.081407.104740.

2. Bebber, D.P., Holmes, T., and Gurr, S.J. (2014). The global spread of crop pests and pathogens. Glob. Ecol. Biogeogr. 23, 1398–1407. 10.1111/geb.12214.

3. Micali, C., Göllner, K., Humphry, M., Consonni, C., and Panstruga, R. (2008). The Powdery Mildew Disease of Arabidopsis: A Paradigm for the Interaction between Plants and Biotrophic Fungi. Arabidopsis Book 6, e0115. 10.1199/tab.0115.

4. Plotnikova, J.M., Reuber, T.L., Ausubel, F.M., and Pfister, D.H. (1998). Powdery mildew pathogenesis of Arabidopsis thaliana. Mycologia 90, 1009–1016. 10.1080/00275514.1998.12026999.

5. Chandran, D., Inada, N., Hather, G., Kleindt, C.K., and Wildermuth, M.C. (2010). Laser microdissection of Arabidopsis cells at the powdery mildew infection site reveals site-specific processes and regulators. Proc. Natl. Acad. Sci. U. S. A. 107, 460–465. 10.1073/pnas.0912492107.

6. Chandran, D., Rickert, J., Cherk, C., Dotson, B.R., and Wildermuth, M.C. (2013). Host cell ploidy underlying the fungal feeding site is a determinant of powdery mildew growth and reproduction. Mol. Plant. Microbe. Interact. 26, 537–545. 10.1094/MPMI-10-12-0254-R.

7. 7. Jaenisch, J., Xue, H., Schläpfer, J., McGarrigle, E.R., Louie, K., Northen, T.R., and Wildermuth, M.C. (2023). Powdery mildew infection induces a non-canonical route to storage lipid formation at the expense of host thylakoid lipids to fuel its spore production. bioRxiv, 2023.12.15.571944. 10.1101/2023.12.15.571944.

8. Jiang, Y., Wang, W., Xie, Q., Liu, N., Liu, L., Wang, D., Zhang, X., Yang, C., Chen, X., Tang, D., et al. (2017). Plants transfer lipids to sustain colonization by mutualistic mycorrhizal and parasitic fungi. Science 356, 1172–1175. 10.1126/science.aam9970.

9. Both, M., Csukai, M., Stumpf, M.P.H., and Spanu, P.D. (2005). Gene expression profiles of Blumeria graminis indicate dynamic changes to primary metabolism during development of an obligate biotrophic pathogen. Plant Cell 17, 2107–2122. 10.1105/tpc.105.032631.

10. McRae, A.G., Taneja, J., Yee, K., Shi, X., Haridas, S., LaButti, K., Singan, V., Grigoriev, I.V., and Wildermuth, M.C. (2023). Spray-induced gene silencing to identify powdery mildew gene targets and processes for powdery mildew control. Mol. Plant Pathol. 24, 1168–1183. 10.1111/mpp.13361.

11. Clark, J.I.M., and Hall, J.L. (1998). Solute transport into healthy and powdery mildew-infected leaves of pea and uptake by powdery mildew mycelium. New Phytol. 140, 261–269. 10.1046/j.1469-8137.1998.00263.x.

12. Sutton, P.N., Henry, M.J., and Hall, J.L. (1999). Glucose, and not sucrose, is transported from wheat to wheat powdery mildew. Planta 208, 426–430. 10.1007/s004250050578.

13. Fotopoulos, V., Gilbert, M.J., Pittman, J.K., Marvier, A.C., Buchanan, A.J., Sauer, N., Hall, J.L., and Williams, L.E. (2003). The monosaccharide transporter gene, AtSTP4, and the cell-wall invertase, Atbetafruct1, are induced in Arabidopsis during infection with the fungal biotroph Erysiphe cichoracearum. Plant Physiol. 132, 821–829. 10.1104/pp.103.021428.

14. Swarbrick, P.J., Schulze-Lefert, P., and Scholes, J.D. (2006). Metabolic consequences of susceptibility and resistance (race-specific and broad-spectrum) in barley leaves challenged with powdery mildew. Plant Cell Environ. 29, 1061–1076. 10.1111/j.1365-3040.2005.01472.x.

15. Wildermuth, M.C. (2010). Modulation of host nuclear ploidy: a common plant biotroph mechanism. Curr. Opin. Plant Biol. 13, 449–458. 10.1016/j.pbi.2010.05.005.

16. Gass, N., Glagotskaia, T., Mellema, S., Stuurman, J., Barone, M., Mandel, T., Roessner-Tunali, U., and Kuhlemeier, C. (2005). Pyruvate decarboxylase provides growing pollen tubes with a competitive advantage in petunia. Plant Cell 17, 2355–2368. 10.1105/tpc.105.033290.

17. Mellema, S., Eichenberger, W., Rawyler, A., Suter, M., Tadege, M., and Kuhlemeier, C. (2002). The ethanolic fermentation pathway supports respiration and lipid biosynthesis in tobacco pollen. Plant J. 30, 329–336. 10.1046/j.1365-313x.2002.01293.x.

18. Wei, Y., Lin, M., Oliver, D.J., and Schnable, P.S. (2009). The roles of aldehyde dehydrogenases (ALDHs) in the PDH bypass of Arabidopsis. BMC Biochem. 10, 7. 10.1186/1471-2091-10-7.

19. Kürsteiner, O., Dupuis, I., and Kuhlemeier, C. (2003). The pyruvate decarboxylase1 gene of Arabidopsis is required during anoxia but not other environmental stresses. Plant Physiol. 132, 968–978. 10.1104/pp.102.016907.

20. Sutak, R., Tachezy, J., Kulda, J., and Hrdý, I. (2004). Pyruvate decarboxylase, the target for omeprazole in metronidazole-resistant and iron-restricted Tritrichomonas foetus. Antimicrob. Agents Chemother. 48, 2185–2189. 10.1128/AAC.48.6.2185-2189.2004.

21. Johnson, D., Weber, D.J., and Hess, W.M. (1976). Lipids from conidia of Erysiphe graminis tritici (powdery mildew). Trans. Br. Mycol. Soc. 66, 35–43. 10.1016/S0007-1536(76)80090-3.

22. Muchembled, J., Sahraoui, A.L.-H., Laruelle, F., Palhol, F., Couturier, D., Grandmougin-Ferjani, A., and Sancholle, M. (2005). Methoxylated fatty acids in Blumeria graminis conidia. Phytochemistry 66, 793–796. 10.1016/j.phytochem.2005.02.011.

23. Tulloch, A.P., and Ledingham, G.A. (1960). The component fatty acids of oils found in spores of plant rusts and other fungi. Can. J. Microbiol. 6, 425–434. 10.1139/m60-048.

24. Jones, L., Riaz, S., Morales-Cruz, A., Amrine, K.C.H., McGuire, B., Gubler, W.D., Walker, M.A., and Cantu, D. (2014). Adaptive genomic structural variation in the grape powdery mildew pathogen, Erysiphe necator. BMC Genomics 15, 1081. 10.1186/1471-2164-15-1081.

25. Fotopoulos, V. (2003). Molecular analysis of nutrient transfer in the host/powdery mildew interaction.

26. Singh, A., and Del Poeta, M. (2011). Lipid signalling in pathogenic fungi. Cell. Microbiol. 13, 177–185. 10.1111/j.1462-5822.2010.01550.x.

27. Tsitsigiannis, D.I., and Keller, N.P. (2007). Oxylipins as developmental and host–fungal communication signals. Trends Microbiol. 15, 109–118. 10.1016/j.tim.2007.01.005.

28. Rangan, P., Maurya, R., and Singh, S. (2022). Can omic tools help generate alternative newer sources of edible seed oil? Plant Direct 6, e399. 10.1002/pld3.399.

29. Vanhercke, T., Dyer, J.M., Mullen, R.T., Kilaru, A., Rahman, M.M., Petrie, J.R., Green, A.G., Yurchenko, O., and Singh, S.P. (2019). Metabolic engineering for enhanced oil in biomass. Prog. Lipid Res. 74, 103–129. 10.1016/j.plipres.2019.02.002.

30. Fell, D.A., Saavedra, E., and Rohwer, J. (2024). 50 years of Metabolic Control Analysis: Its past and current influence in the biological sciences. Biosystems. 235, 105086. 10.1016/j.biosystems.2023.105086.

31. Kacser, H., and Burns, J.A. (1973). The control of flux. Symp. Soc. Exp. Biol. 27, 65–104.

32. Fell, D.A., Taylor, D.C., Weselake, R.J., and Harwood, J.L. (2023). Metabolic Control Analysis of triacylglycerol accumulation in oilseed rape. Biosystems. 227-228, 104905. 10.1016/j.biosystems.2023.104905.

33. Hirani, T.A., Tovar-Méndez, A., Miernyk, J.A., and Randall, D.D. (2011). Asp295 stabilizes the active-site loop structure of pyruvate dehydrogenase, facilitating phosphorylation of ser292 by pyruvate dehydrogenase-kinase. Enzyme Res. 2011, 939068. 10.4061/2011/939068.

34. Joshi, J., Folz, J.S., Gregory, J.F., III, McCarty, D.R., Fiehn, O., and Hanson, A.D. (2019). Rethinking the PDH bypass and GABA shunt as thiamin-deficiency workarounds. Plant Physiol. 181, 389–393.

35. Ye, S. (2009). Identification of pyruvate decarboxylase/indole pyruvate decarboxylase gene family members from Arabidopsis thaliana.

36. Vromman, F., and Subtil, A. (2014). Exploitation of host lipids by bacteria. Curr. Opin. Microbiol. 17, 38–45. 10.1016/j.mib.2013.11.003.

37. Caffaro, C.E., and Boothroyd, J.C. (2011). Evidence for host cells as the major contributor of lipids in the intravacuolar network of Toxoplasma-infected cells. Eukaryot. Cell 10, 1095–1099. 10.1128/EC.00002-11.

38. Itoe, M.A., Sampaio, J.L., Cabal, G.G., Real, E., Zuzarte-Luis, V., March, S., Bhatia, S.N., Frischknecht, F., Thiele, C., Shevchenko, A., et al. (2014). Host cell phosphatidylcholine is a key mediator of malaria parasite survival during liver stage infection. Cell Host Microbe 16, 778–786. 10.1016/j.chom.2014.11.006.

39. Yu, Y., Maguire, T.G., and Alwine, J.C. (2012). Human cytomegalovirus infection induces adipocyte-like lipogenesis through activation of sterol regulatory element binding protein 1. J. Virol. 86, 2942–2949. 10.1128/JVI.06467-11.

40. Roingeard, P., and Melo, R.C.N. (2017). Lipid droplet hijacking by intracellular pathogens. Cell. Microbiol. 19. 10.1111/cmi.12688.

41. Nolan, S.J., Romano, J.D., and Coppens, I. (2017). Host lipid droplets: An important source of lipids salvaged by the intracellular parasite Toxoplasma gondii. PLoS Pathog. 13, e1006362. 10.1371/journal.ppat.1006362.

42. Hüsler, D., Stauffer, P., and Hilbi, H. (2023). Tapping lipid droplets: A rich fat diet of intracellular bacterial pathogens. Mol. Microbiol. 120, 194–209. 10.1111/mmi.15120.

43. Wilburn, K.M., Fieweger, R.A., and VanderVen, B.C. (2018). Cholesterol and fatty acids grease the wheels of Mycobacterium tuberculosis pathogenesis. Pathog. Dis. 76. 10.1093/femspd/fty021.

44. 44. Brancucci, N.M.B., Gerdt, J.P., Wang, C., De Niz, M., Philip, N., Adapa, S.R., Zhang, M., Hitz, E., Niederwieser, I., Boltryk, S.D., et al. (2017). Lysophosphatidylcholine Regulates Sexual Stage Differentiation in the Human Malaria Parasite Plasmodium falciparum. Cell 171, 1532–1544.e15. 10.1016/j.cell.2017.10.020.

45. MacLean, A.M., Bravo, A., and Harrison, M.J. (2017). Plant Signaling and Metabolic Pathways Enabling Arbuscular Mycorrhizal Symbiosis. Plant Cell 29, 2319–2335. 10.1105/tpc.17.00555.

46. Kameoka, H., and Gutjahr, C. (2022). Functions of Lipids in Development and Reproduction of Arbuscular Mycorrhizal Fungi. Plant Cell Physiol. 63, 1356–1365. 10.1093/pcp/pcac113.

47. Keymer, A., Pimprikar, P., Wewer, V., Huber, C., Brands, M., Bucerius, S.L., Delaux, P.-M., Klingl, V., Röpenack-Lahaye, E. von, Wang, T.L., et al. (2017). Lipid transfer from plants to arbuscular mycorrhiza fungi. Elife 6. 10.7554/eLife.29107.

48. Bravo, A., Brands, M., Wewer, V., Dörmann, P., and Harrison, M.J. (2017). Arbuscular mycorrhiza-specific enzymes FatM and RAM2 fine-tune lipid biosynthesis to promote development of arbuscular mycorrhiza. New Phytol. 214, 1631–1645. 10.1111/nph.14533.

49. 49. Luginbuehl, L.H., Menard, G.N., Kurup, S., Van Erp, H., Radhakrishnan, G.V., Breakspear, A., Oldroyd, G.E.D., and Eastmond, P.J. (2017). Fatty acids in arbuscular mycorrhizal fungi are synthesized by the host plant. Science 356, 1175–1178. 10.1126/science.aan0081.

50. Schwelm, A., Fogelqvist, J., Knaust, A., Jülke, S., Lilja, T., Bonilla-Rosso, G., Karlsson, M., Shevchenko, A., Dhandapani, V., Choi, S.R., et al. (2015). The Plasmodiophora brassicae genome reveals insights in its life cycle and ancestry of chitin synthases. Sci. Rep. 5, 1–12. 10.1038/srep11153.

51. Liang, P., Liu, S., Xu, F., Jiang, S., Yan, J., He, Q., Liu, W., Lin, C., Zheng, F., Wang, X., et al. (2018). Powdery Mildews Are Characterized by Contracted Carbohydrate Metabolism and Diverse Effectors to Adapt to Obligate Biotrophic Lifestyle. Front. Microbiol. 9, 3160. 10.3389/fmicb.2018.03160.

52. Gomez, S.K., Javot, H., Deewatthanawong, P., Torres-Jerez, I., Tang, Y., Blancaflor, E.B., Udvardi, M.K., and Harrison, M.J. (2009). Medicago truncatula and Glomus intraradices gene expression in cortical cells harboring arbuscules in the arbuscular mycorrhizal symbiosis. BMC Plant Biol. 9, 10. 10.1186/1471-2229-9-10.

53. Gravot, A., Richard, G., Lime, T., Lemarié, S., Jubault, M., Lariagon, C., Lemoine, J., Vicente, J., Robert-Seilaniantz, A., Holdsworth, M.J., et al. (2016). Hypoxia response in Arabidopsis roots infected by Plasmodiophora brassicae supports the development of clubroot. BMC Plant Biol. 16, 251. 10.1186/s12870-016-0941-y.

54. Shiba, Y., Paradise, E.M., Kirby, J., Ro, D.-K., and Keasling, J.D. (2007). Engineering of the pyruvate dehydrogenase bypass in Saccharomyces cerevisiae for high-level production of isoprenoids. Metab. Eng. 9, 160–168. 10.1016/j.ymben.2006.10.005.

55. Chen, Y., Daviet, L., Schalk, M., Siewers, V., and Nielsen, J. (2013). Establishing a platform cell factory through engineering of yeast acetyl-CoA metabolism. Metab. Eng. 15, 48–54. 10.1016/j.ymben.2012.11.002.

56. Koivuranta, K., Castillo, S., Jouhten, P., Ruohonen, L., Penttilä, M., and Wiebe, M.G. (2018). Enhanced Triacylglycerol Production With Genetically Modified Trichosporon oleaginosus. Front. Microbiol. 9, 1337. 10.3389/fmicb.2018.01337.

57. Liu, X., Cooper, D.E., Cluntun, A.A., Warmoes, M.O., Zhao, S., Reid, M.A., Liu, J., Lund, P.J., Lopes, M., Garcia, B.A., et al. (2018). Acetate Production from Glucose and Coupling to Mitochondrial Metabolism in Mammals. Cell 175, 502–513.e13. 10.1016/j.cell.2018.08.040.

58. Prochownik, E.V., and Wang, H. (2021). The Metabolic Fates of Pyruvate in Normal and Neoplastic Cells. Cells 10. 10.3390/cells10040762.

59. Vysochan, A., Sengupta, A., Weljie, A.M., Alwine, J.C., and Yu, Y. (2017). ACSS2-mediated acetyl-CoA synthesis from acetate is necessary for human cytomegalovirus infection. Proc. Natl. Acad. Sci. U. S. A. 114, E1528–E1535. 10.1073/pnas.1614268114.

60. Austin, L.S., Kaushansky, A., and Kappe, S.H.I. (2014). Susceptibility to Plasmodium liver stage infection is altered by hepatocyte polyploidy. Cell. Microbiol. 16, 784–795. 10.1111/cmi.12282.

61. Anatskaya, O.V., and Vinogradov, A.E. (2007). Genome multiplication as adaptation to tissue survival: evidence from gene expression in mammalian heart and liver. Genomics 89, 70–80. 10.1016/j.ygeno.2006.08.014.

62. Shi, Q., Xue, C., Zeng, Y., Gu, X., Wang, J., and Li, L. (2023). A novel prognostic model for hepatocellular carcinoma based on pyruvate metabolism-related genes. Sci. Rep. 13, 9780. 10.1038/s41598-023-37000-8.

