## Supplemental Information for "Altered host pyruvate metabolism fuels and regulates fungal asexual reproduction"

### SUPPLEMENTARY INFORMATION OVERVIEW

- **Supplemental Document 1:** Supplemental Overview, Figures S1-S4, Tables S1-S2.
- **Excel File S1**
- **Excel Workbook File S2**

#### Supplemental Figures

**Figure S1.** Characterization of Arabidopsis *pdc2-1* T-DNA insertion mutant, supports Figure 2 and STAR Methods.

**Figure S2.** Representative images of *aldh2B4,B7* double mutant compared with WT plants infected with powdery mildew, supports Figure 2.

**Figure S3**. Spore TGs contain C20-C24 acyl chains (MS2 analysis), supports Figure 3.

**Figure S4**. Representative TG signatures for spore and washed leaf lipids for detached leaves supplied with ethanol vs. $^{13}C$-ethanol, supports Figure 3.

#### Supplemental Tables and Excel Files

**Table S1:** Complementation of *pdc2* mutant, supports Figure 2.

**Table S2:** Primers Used, supports STAR Methods.

**Excel File S1.** Reciprocal tBLASTn results show no PDC or ADH candidate in *G. orontii MGH1*, supports STAR Methods. Provided as a separate file.

**Excel File S2.** TG identification, relative abundance and $^{13}C$ enrichment results, Supports Figure 3. Provided as a separate file.

### Figure S1. Characterization of Arabidopsis *pdc2-1* T-DNA insertion mutant, supports Figure 2 and STAR Methods.

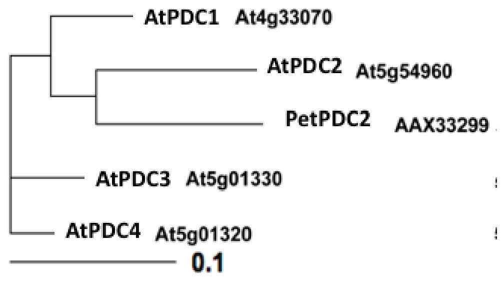


**A. ClustalW analysis (Blosum62 matrix) of Arabidopsis PDC proteins compared with Petunia PDC2 (PetPDC2).** PetPDC2 plays a role in the PDH bypass during pollen development (Gass et al. (2005) *Plant Cell* **17**:2355-68). Generated using TreeViewX.

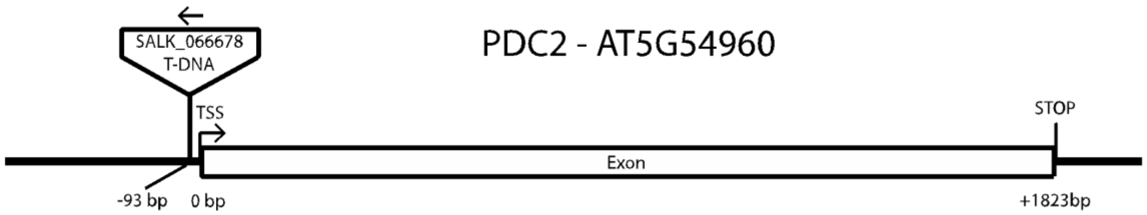


**B. Gene structure of *AtPDC2* and location of T-DNA insertion in *pdc2-1* (SALK_066678).** *pdc2-1* mutant seed were obtained from the Arabidopsis Biological Resource Center and confirmed to be homozygous with T-DNA insertion site shown above. Primers used for genotyping are included in Table S2.

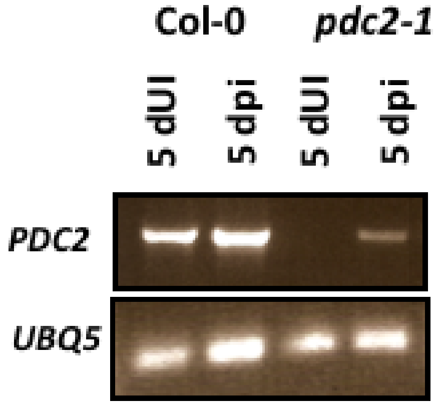


**C. *pdc2-1* exhibits dramatically reduced expression in powdery mildew-infected leaves.** Expression of *PDC2* and housekeeping gene *Ubiquitin5* in mature fully expanded leaves at 5 days post infection (dpi) with powdery mildew compared with uninfected plants (dUI) grown in parallel. Primers for amplification of transcript by RT-PCR are in Table S2.

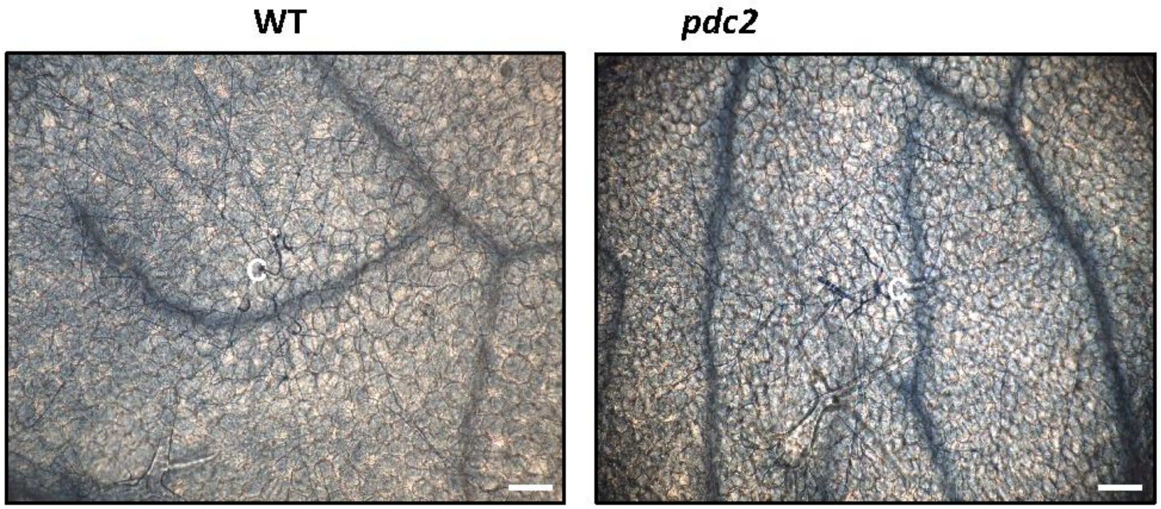


**D. Cell death is not observed in *pdc2* mutant.** Representative image of mesophyll cells underlying fungal colony at 5 dpi for WT and *pdc2-1* mutant, stained with trypan blue to observe cell death microscopically. See Methods for details. C= initial germinated conidia; Bar= 100 um. Independent experiments gave similar results.

**Figure S2. Representative images of *aldh2B4,B7* compared with WT plants infected with powdery mildew, supports Figure 2.**

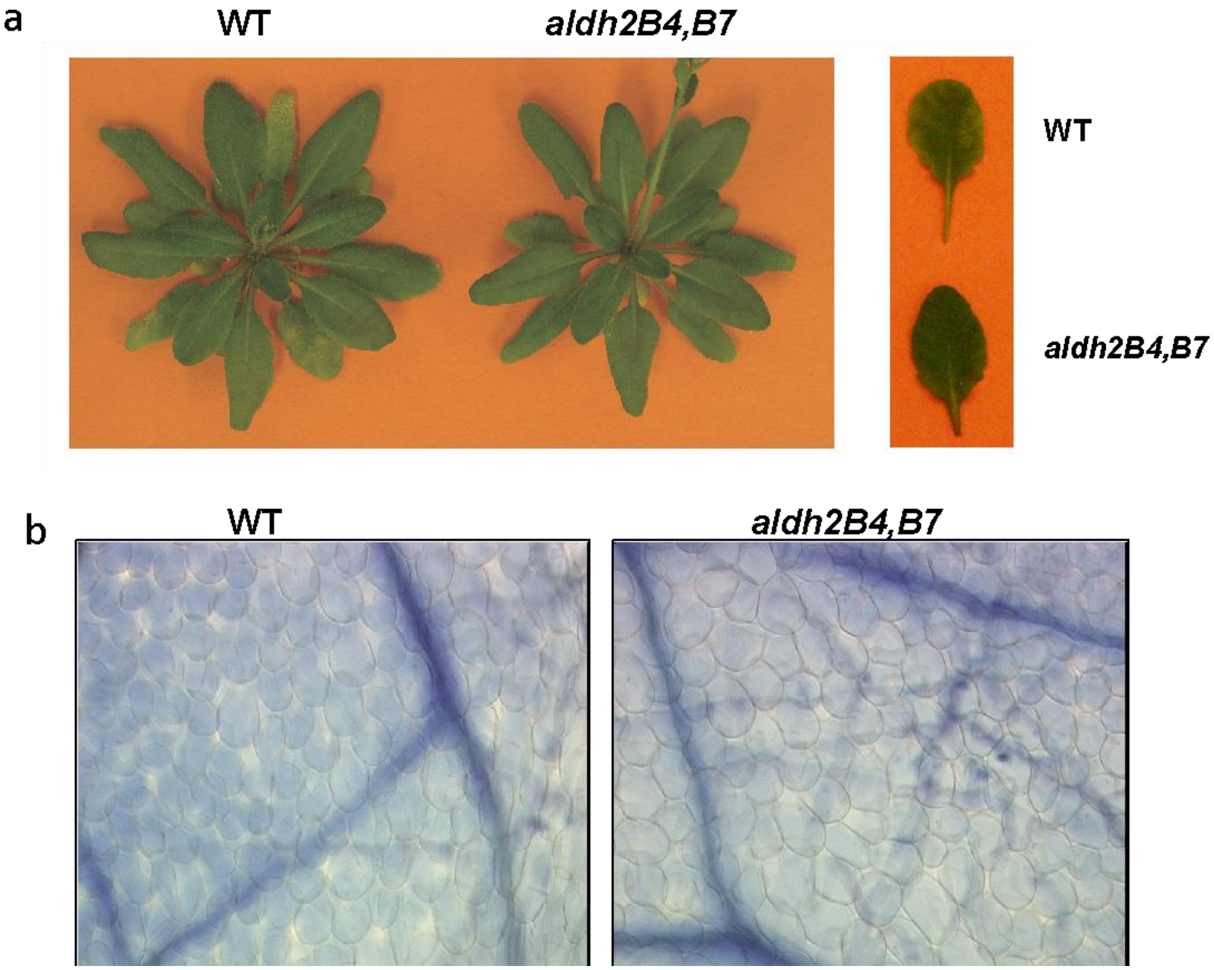


Representative image of powdery mildew growth on WT and *aldh2B4aldh2B7* double mutant plants **(A)** at 11 dpi, and **(B)** mesophyll cells underlying fungal colony at 5 dpi, stained with trypan blue to observe cell death microscopically at 20x. See Methods for details.

**Figure S3. Representative MS2 spectra showing spore TGs with C20-C24 acyl chains, supports Figure 3.**

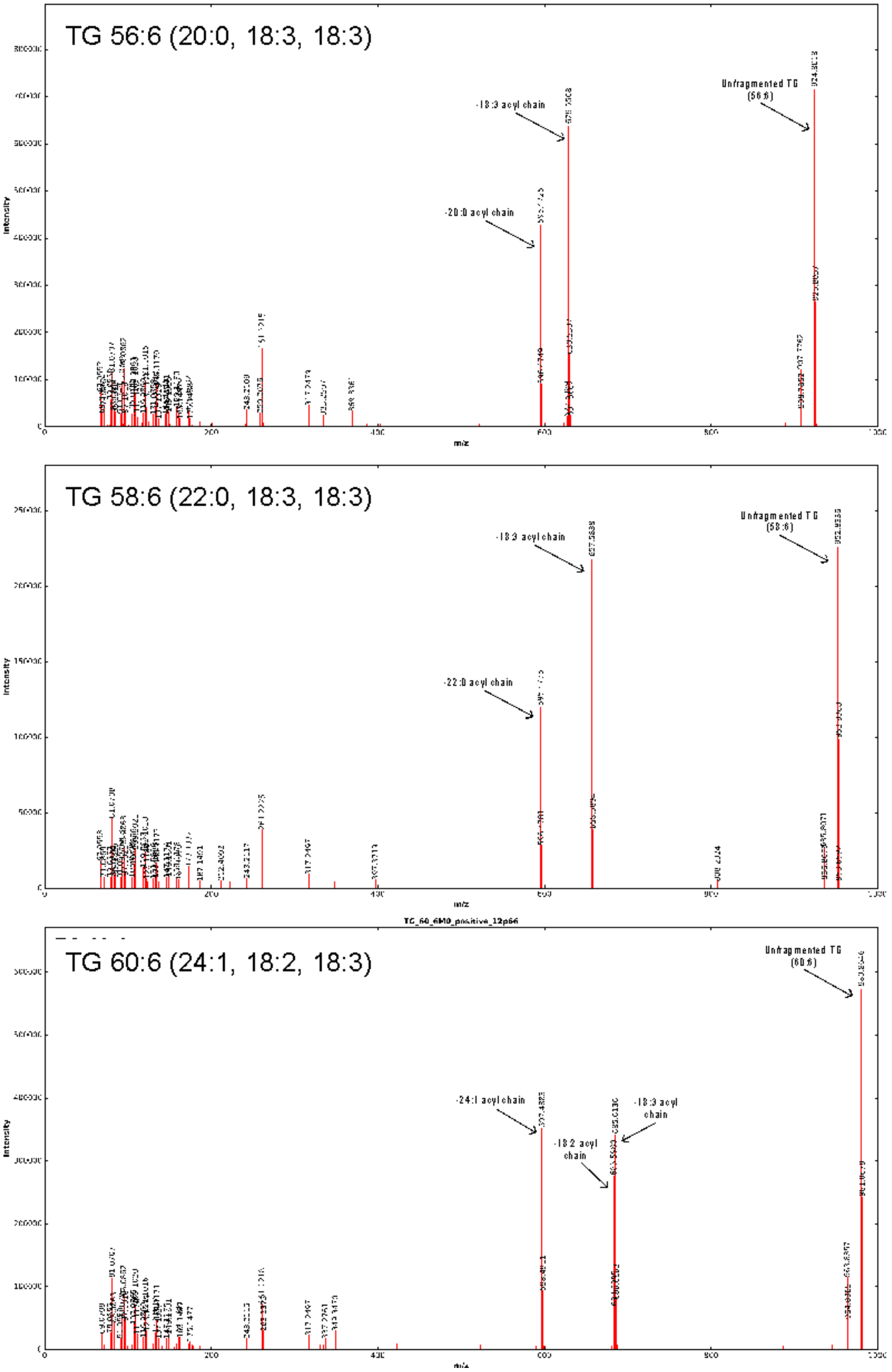


MS2 spectra (positive ion mode) for selected TGs is shown. TG acyl chain assignments utilized theoretical m/z for $M_0$ and neutral loss calculations for dominant fragments in MS2.

**Figure S4. Representative TG signatures for spore and washed leaf lipids for detached leaves supplied with ethanol vs. $^{13}$C-ethanol, supports Figure 3.**

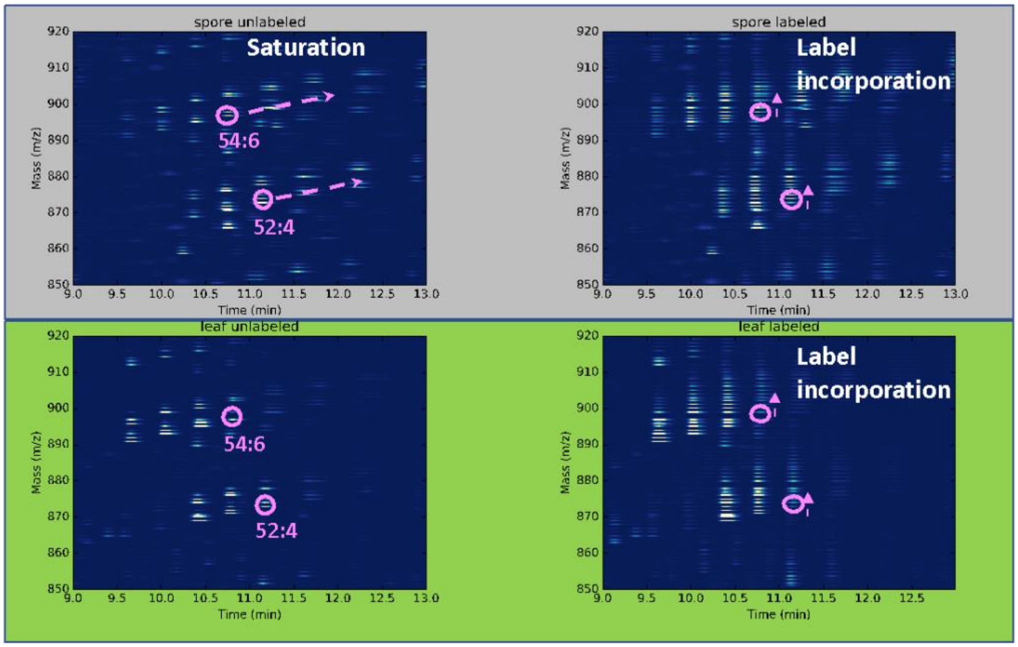


Representative 2D heat maps are shown for spores (top) and washed leaves (bottom) from agar plates with ethanol (unlabeled) versus 13C-ethanol (labeled). Retention time and m/z shown profile TGs with TG saturation (dotted arrow) in spores vs. leaves illustrated for TG 54:6 and 52:4. Multiple label incorporation is illustrated for both spore and leaf TG 54:6 and 52:4 (solid arrow).

#### Table S1. Complementation of *pdc2*, supports Figure 2.

| **Genotype** | **cp/colony** | **STDEV** | **ttest p-value** | |
|---|---|---|---|---|
| WT Col-0 | 5.79 | 3.3 | w/Col-0 | w/pdc2 |
| *pdc2* | 3.89 | 2.96 | 7.13E-05 | |
| *PDC2/pdc2* | 5.19 | 4.11 | 2.81E-01 | 1.58E-02 |

Complementation of *pdc2-1* with *PDC2* restored powdery mildew asexual reproduction to WT levels. Powdery mildew assay and creation of stable complemented homozygous *PDC2/pdc2* transgenic line is described in Methods. cp=conidiophore.

### Table S2 Primers Used, Supports STAR Methods.

| Name | Sequence (5' to 3') | Purpose |
|---|---|---|
| ***PDC2pro::PDC2-eGFP* construct in pCAMBIA 1300** | | |
| PDC2HindIII-F | GCGCAAGCTTAAAGAGATTGTGGGAGATATGTGATTG | PDC2 |
| PstIPDC2-R | AACTGCAGCTGCGGATTTGGGGGACGAC | PDC2 |
| PDC2 pro PstI-R | AAACTGCAGTTGAAATATGAGATTAGTGG | PDC2 promoter |
| eGFP PstI-F | AAACTGCAGATGGTGAGCAAGGGCGAGGAGC | eGFP |
| eGFP SalI-R | ACGCGTCGACTTACTTGTACAGCTCGTCCATG | eGFP |
| SalI35S term-F | ACGCGTCGACAGCTCGAATTCGGTACGCTGAAATCAC | 35S terminator |
| SacI35S term-R | GCGCGAGCTCGGGTTTTCCCAGTCACGACGTTG | 35S terminator |
| *PDC2 F* | AGTCCACTTCACTCCACCAC | RT-PCR |
| *PDC2 R* | CACTAACTCCTCCTCGCATC | RT-PCR |
| **Genotyping mutants** | | |
| *pdc2-1* F | GAGGAATCGCCGGTAAATTAC | *pdc2-1* (SALK_066678C) |
| *pdc2-1* R | ATTGGAGATTGGGGATCTACG | *pdc2-1* |

| | | |
|---|---|---|
| *aldhb2b4-1* F | GTTGGTCCTGCTCTTGCTTGTGGTAA | *aldhb2b4-1* |
| *aldhb2b4-1* R | TCGTTCGCCCTCTTTATCACCTCATC | aldhb2b4-1 |
| *aldhb2b7-1* F | TTGAGACTTGGGATAATGGGAAACCT | *aldhb2b7-1* |
| *aldhb2b7-1* R | AAGAAAACTGTGACGGTAATAATCGG | *aldhb2b7-1* |
| *aldh2b4-2 F* | ATTCAAAGTACGGCAACACAAACCAAGAG | *aldhb2b4-2 (SALK_078568)* |
| *aldh2b4-2 R* | TTACCACAAGCAAGAGCAGGACCAAC | *aldhb2b4-2* |
| *adh1-1* F | TCTCTCTGAAGGCTGGAGATG | *adh1-1 (SALK_066824)* |
| *aldh1-1* R | AAACCCAAAACTGACATTCCC | *adh1-1* |
| *LBb1.3* | ATTTTGCCGATTTCGGAAC | *Left border of T-DNA (SALK)* |

Primers for genotyping *aldh* mutants were previously described in Wei et al. (2009) *BMC Biochemistry* **10**:7. F= forward, R= reverse.
